# Sucrase isomaltase dysfunction influences dietary sucrose intake and preference

**DOI:** 10.1101/2024.04.18.590042

**Authors:** Peter Aldiss, Leire Torices, Stina Ramne, Marit Eika Jørgensen, Sucrase-Isomaltase Working Group, Mauro D’Amato, Mette K Andersen

## Abstract

**Objective:** To characterise the role of Sucrase-isomaltase (SI) in regulating dietary behaviours, such as sweet preference and food liking in Si knockout (Sis-KO) mice and in population-based cohorts from Greenland and the UK.

**Design:** We profiled the appetitive and post-ingestive response to dietary carbohydrates in SI knockout (Sis-KO) mice. Alongside this, we conducted detailed dietary analysis of 45 foods in two Greenlandic population-based cohorts (IHIT, n=2778 and 68 foods, and B2018, n=2203 and 45 foods) with the presence of a common (allele frequency = 14.3%) SI Loss of function (LoF) variant, c.273-274delAG. Finally, we explored the association between SI hypomorphic variants, liking of 140 foods, and sucrose content using data from 134,766 UKBB participants with exome sequencing and questionnaire data available.

**Results:** Sucrose naïve Sis-KO mice had a significantly reduced intake of dietary sucrose, and preference for 10% liquid sucrose, in two-bottle preference studies. Mechanistically, oral administration of the short-chain fatty acid acetate reduced sucrose-preference in wild-type mice. In Greenlandic LoF homozygous carriers we show that the previously reported reduction in sugar intake may primarily be explained by a lower intake of cake and pastries, and of candy and chocolate and that added sugar is the main factor explaining these associations. In the UKBB, a negative association with “cake icing”, the food with the highest sucrose content per 100g, was detected in SI hypomorphic carriers, as well as in sensitivity analyses conducted only including carriers of known CSID LoF variants. Further, a negative linear relationship was also observed between the effect estimates of hypomorphic SI variants on food liking and the estimated sucrose content per 100g of 88 sucrose-containing foods, indicating that food dislike in SI carriers correlates with the amount of sucrose in food.

**Conclusion:** Collectively, we demonstrated that genetic variation in the SI gene is associated with significant changes in sucrose preference, characterised by a rapid avoidance of dietary sucrose in Sis-KO mice, as well as lower consumption and increased disliking of sucrose rich foods in Greenlanders and Europeans, respectively. This work demonstrates that genetic variation in the SI gene may impact physiology beyond the gastrointestinal tract and suggest the possibility to target SI to reduce the preference, and intake, of dietary sucrose with implications for digestive and metabolic health.

## MAIN TEXT

Carbohydrates constitute a large proportion of the modern diet and play a major role in both health and disease. For instance, excess intake of dietary sugar is established to be a contributor to obesity and type 2 diabetes (T2D) whilst a subset of carbohydrates poorly absorbed in the small intestine are recognized triggers of, and therapeutically targeted in irritable bowel syndrome (IBS) [1]. Homozygous or compound heterozygous loss-of-function (LoF) variants in the sucrase-isomaltase (*SI*) gene drive congenital SI deficiency (CSID), whilst dysfunctional (hypomorphic) variants have been reported to increase the risk of IBS and to affect the response to carbohydrate-restricted diets [2-4]. In Greenlandic adults, we recently reported that c.273-274delAG *SI* LoF homozygotes exhibit lower BMI, improved metabolic health, a lower intake of added sugar, and no significant increase in selfreported gastrointestinal symptoLoF homozygotes exhibitms [5]. However, in the electronic health records we observed an increased number of contacts and gastrointestinal related diagnostic procedures in carriers compared to non-carriers [6]. Here, we expand these findings to characterise the role of SI in regulating dietary behaviours, such as sweet preference and food liking in *Si* knockout (Sis-KO) mice and in population-based cohorts from Greenland and the UK.

First, we showed that sucrose naïve Sis-KO mice had a significantly reduced preference for liquid 10% sucrose during two-bottle preference studies with water as the other choice, and lower intake of dietary 17% sucrose (Fig. 1A-B), consuming ∼66% less of each. This demonstrates a striking ability for SI loss of function to regulate sucrose preference, and intake, whilst simultaneously increasing the preference for other carbohydrates and non-caloric sweeteners (Fig. S1A-C). Using metabolic cages, we demonstrated that sucrose avoidance developed rapidly within the first hour of access in sucrose naïve Sis-KO mice (Fig. 1C-D). Then, we showed that an oral sucrose gavage in Sis-KO mice was associated with alterations in the concentration of the post-ingestive appetitive peptides glucose-dependent insulinotropic polypeptide (GIP) and glucagon-like peptide 1 (GLP-1), which may alter appetite and satiety (Fig. 1E-F). To elucidate the mechanism driving the reduction in sucrose preference we demonstrated that oral administration of the short-chain fatty acid acetate, which is rapidly increased in plasma of Sis-KO mice following sucrose ingestion and is elevated in the plasma of human c.273-274delAG *SI* LoF homozygotes carriers [5], reduced the sucrose-preference in wild-type mice (Fig. 1G). This suggests that gut-derived acetate is a potential mediator, not only of the healthier metabolic phenotype, but also the rapid sucrose avoidance.

**Figure 1.**
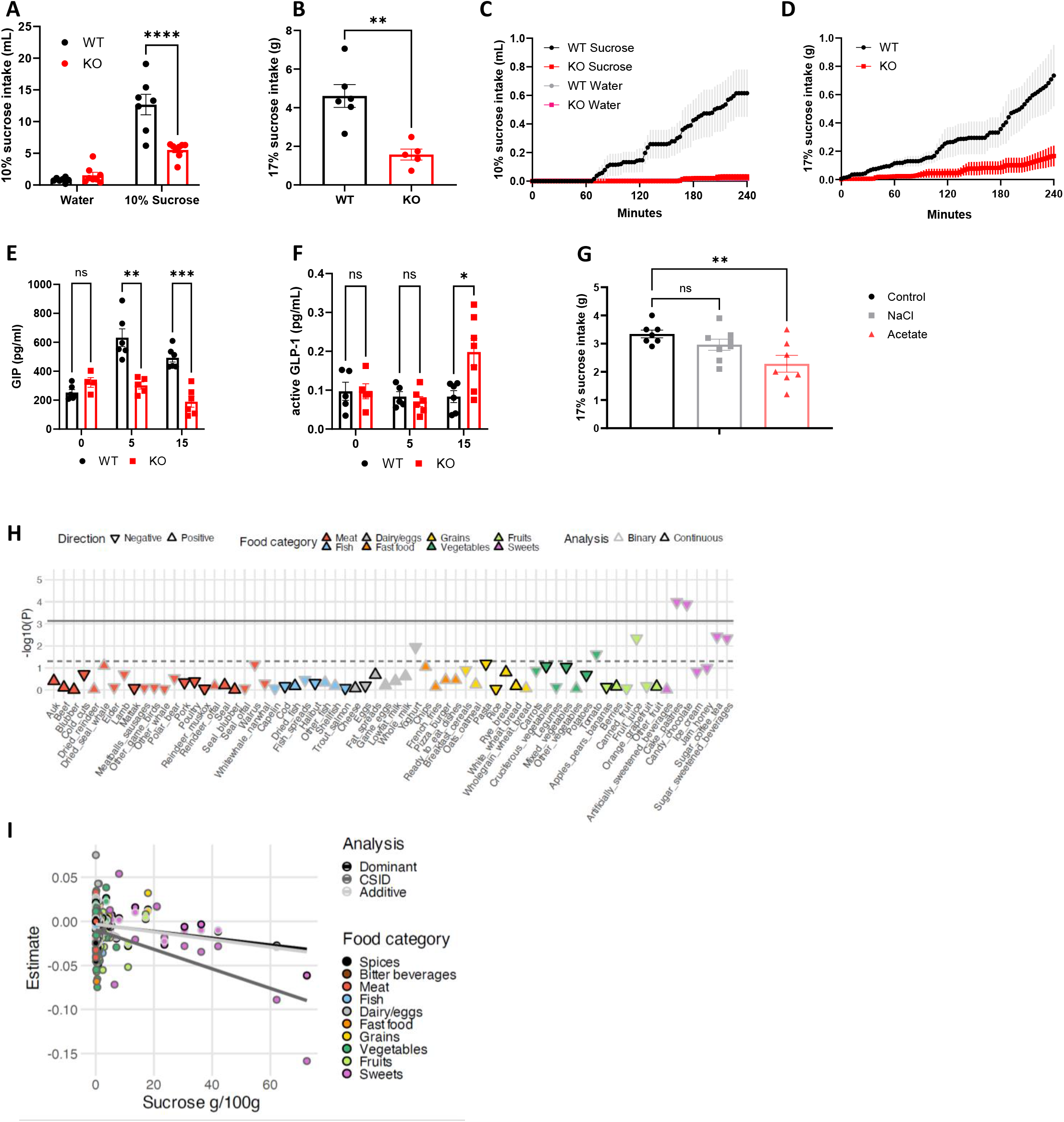
Wild-type and Sis-knockout mouse preference for 10% liquid sucrose (A), intake of 17% dietary sucrose (B) and first 6 hours intake of 10% liquid sucrose (C) and 17% dietary sucrose (D). Post-ingestive appetitive hormones glucose-dependent insulinotropic polypeptide (GIP, E) and active glucagon-like peptide 1 (GLP-1, F) following an oral gavage of sucrose (3g/kg). Intake of dietary sucrose following supplementation of the drinking water with either sodium chloride (sodium control) or sodium acetate (G). *p<0.05, **p<0.01, ***p<0.001, ****p<0.00001. Analysis was carried out using a two-way ANOVA (A, E, F), t-test (B) or one-way ANOVA (G). Analysis of associations between the Greenlandic c.273-274delAG homozygous carriers and selfreported intake of 68 food items (H) using a recessive model in the IHIT cohort (n=2778). The solid horizontal line represents the Bonferroni adjusted significance cutoff (P-value of 0.05/68) and the dashed horizontal line represents a P-value of 0.05. Association analysis between sucrose content of food items and effect sizes from ordered logistic regressions (I) under three models. ‘Dominant’ model where 0 = no variant, 1 = one or more variants, n=10577 carriers; ‘Additive’ model where 0 = no variant, 1 = one variant, 2 = two or more variants, n=10107 single carriers and 470 multiple-carriers; ‘CSID’, sensitivity analyses where 0 = no variants, 1 = one or more variants previously identified in CSID patients and confirmed as loss-of-function alleles in in vitro experiments, n= 124189 non-carriers and 2381 carriers.

We then explored differences in human dietary patterns, by assessing food intake in two Greenlandic population-based cohorts as well as food liking in UK Biobank (UKBB). In the Greenlandic population we took advantage of the presence of a relatively common (allele frequency = 14.3%) SI LoF variant, c.273-274delAG, which we studied in relation to data from food intake questionnaires from two population-based cohorts (IHIT, n=2778 and 68 foods, and B2018, n=2203 and 45 foods). Here we showed that the reduction in added sugar intake, previously reported in Greenlandic LoF homozygous carriers [5] (-28.55[7.92] g/day, p=2.8x10^-7^), may primarily be explained by a lower intake of cake and pastries, and of candy and chocolate (Fig. 1H and S1D). To investigate the nuances of sugar intake (Fig. S1E), combined analyses between the two Greenlandic cohorts demonstrated that c.273-274delAG homozygotes showed lower intake of foods high in added sugar (βSD[SE] =-0.51 [0.1], P=3.9x10^-7^), sugar and fat (βSD[SE] =-0.35[0.1], P=9.1x10^-4^), and of sugar beverages (βSD[SE]=-0.32 [0.1], P=1.1x10^-3^), while total intake of total sugar-rich foods did not differ from the rest of the population. This aligns with the individual food item analyses and points to added sugar, which is generally sucrose, as the main factor explaining these associations.

Finally, we explored the association between *SI* hypomorphic variants [7-9] and liking of 140 foods using data from 134,766 UKBB participants with exome sequencing and questionnaire data available. When 140 food liking traits were individually tested (using ordered logistic regression adjusted for age, sex and top 10 principal components, see Supplementary Methods), a negative association with “cake icing”, the food with the highest sucrose content per 100g, was detected in *SI* hypomorphic carriers (β[SE] = -0.06[0.02], P_adjusted_= 0.04 under an additive model), as well as in sensitivity analyses conducted only including carriers of known CSID LoF variants (β[SE] = -0.16 [0.04], P_adjusted_= 0.005) (Fig. S1F). Further, a negative linear relationship was also observed between the effect estimates of hypomorphic *SI* variants on food liking and the estimated sucrose content per 100g of 88 sucrose-containing foods (best p = 0.0001 for CSID LoF variants, Fig. 1I), indicating that food dislike in SI carriers correlates with the amount of sucrose in food.

Collectively, we demonstrated that genetic variation in the *SI* gene is associated with significant changes in sucrose preference, characterised by a rapid avoidance of dietary sucrose in Sis-KO mice, as well as lower consumption and increased disliking of sucrose rich foods in Greenlanders and Europeans, respectively. In mice, we demonstrated that the avoidance of sweet foods is sucrose-specific rather than broadly directed towards carbohydrates or sweetness and that a loss of function in the *SI* gene can increase preference for other carbohydrates and non-nutritive sweeteners. Mechanistically, we demonstrated that loss-of-function in the *SI* gene is associated with alterations in secretion of the appetitive peptides GLP-1 and GIP following sucrose ingestion in mice. Further, the reduction in sucrose preference in mice may be driven by gut derived production of acetate from undigested sucrose that, which could have broad effects on body weight, peripheral insulin sensitivity, and appetite regulation through the central nervous system. This work demonstrates that genetic variation in the *SI* gene may impact physiology beyond the gastrointestinal tract. These findings are of potential impact at the population level and suggest the possibility to target *SI* to reduce the preference, and intake, of dietary sucrose with implications for digestive and metabolic health.

## CRediT

**Conceptualization:** PA, MPG, MDA, MKA, IM, AA, TH; **Data curation:** PA, CEB, FB, LT, MEJ, FFVS, NKS, NG, PB, IM, AA; **Formal analysis:** PA, SR, FFVS, FB, LT; **Funding acquisition:** PA, MPG, MDA, TH; **Investigation:** PA, SR, MKA, CEB, FB, LT HB; **Methodology:** PA, MPG, SR, MKA, CEB, FB, LT; **Project administration:** PA, MPG, TH, MKA, MDA; **Resources:** MPG, TH, MDA, JT, TG**; Supervision:** PA, MPG, MEJ, MKA, TH, MDA; **Visualization:** PA, SR, LT; **Writing - original draft:** PA, SR, MKA, LT, MDA; **Writing - review** & editing: PA, CEB, FB, LT, MDA, SR, MEJ, MPG, MKA, FFVS, NKS, NG, HB, JTT, TG, PB, IM, AA, TH

## Acknowledgements

**Sucrase-isomaltase working group:** Ferdinando Bonfiglio^1^, Hayley Burm^5^, Cristina Esteban Blanco^2^, Frederik Filip Vinggaard Stæger^3^, Ninna Karsbæk Senftleber^4^, Jonas T. Treebak^5^, Niels Grarup^5^, Peter Bjerregaard^6^, Trisha J. Grevengoed^7^, Matthew P. Gillum^5^, Ida Moltke^3^, Anders Albrechtsen^3^, Torben Hansen^5^

1. CEINGE Biotecnologie Avanzate s.c.ar.l., Naples, Italy; Department of Chemical, Materials and Production Engineering, University of Naples Federico II, Naples, Italy.
2. Gastrointestinal Genetics Lab, CIC bioGUNE – BRTA, Derio, Spain.
3. Department of Biology, University of Copenhagen, Copenhagen, Denmark.
4. Clinical and Translational Research, Copenhagen University Hospital, Steno Diabetes Center Copenhagen, Herlev, Denmark; Steno Diabetes Center Greenland, Queen Ingrid’s Hospital, Nuuk, Greenland.
5. Novo Nordisk Foundation Center for Basic Metabolic Research, Faculty of Health and Medical Sciences, University of Copenhagen, 2200 Copenhagen, Denmark.
6. Centre for Public Health in Greenland, National Institute of Public Health, University of Southern Denmark, Copenhagen, Denmark.
7. Department of Biomedical Sciences, University of Copenhagen, Copenhagen, Denmark

## Supplementary Figure 1

Wild-type and Sis-knockout mouse preference for 10% liquid Glucose (A), Maltodextrin (B) and 0.8% Sucralose (C), *p<0.05, **p<0.01, ****p<0.00001.

Dietary patterns in Greenlandic c.273-274delAG homozygous carriers in the B2018 cohort (D, n=2203)

Merged food group analysis of Greenlandic c.273-274delAG homozygous carriers (E, n=3388).

Analysis of food liking traits from the UK Biobank (F) under three models. ‘Dominant’ model where 0 = no variant, 1 = one or more variants, n=10577 carriers; ‘Additive’ model where 0 = no variant, 1 = one variant, 2 = two or more variants, n=10107 single carriers and 470 multiple-carriers; ‘CSID’, sensitivity analyses where 0 = no variants, 1 = one or more variants previously identified in CSID patients and confirmed as loss-of-function alleles in *in vitro* experiments, n= 124189 non-carriers and 2381 carriers.

