## Supplementary figures and images for "Sucrase isomaltase dysfunction influences dietary sucrose intake and preference"

### Supp. Fig. 1

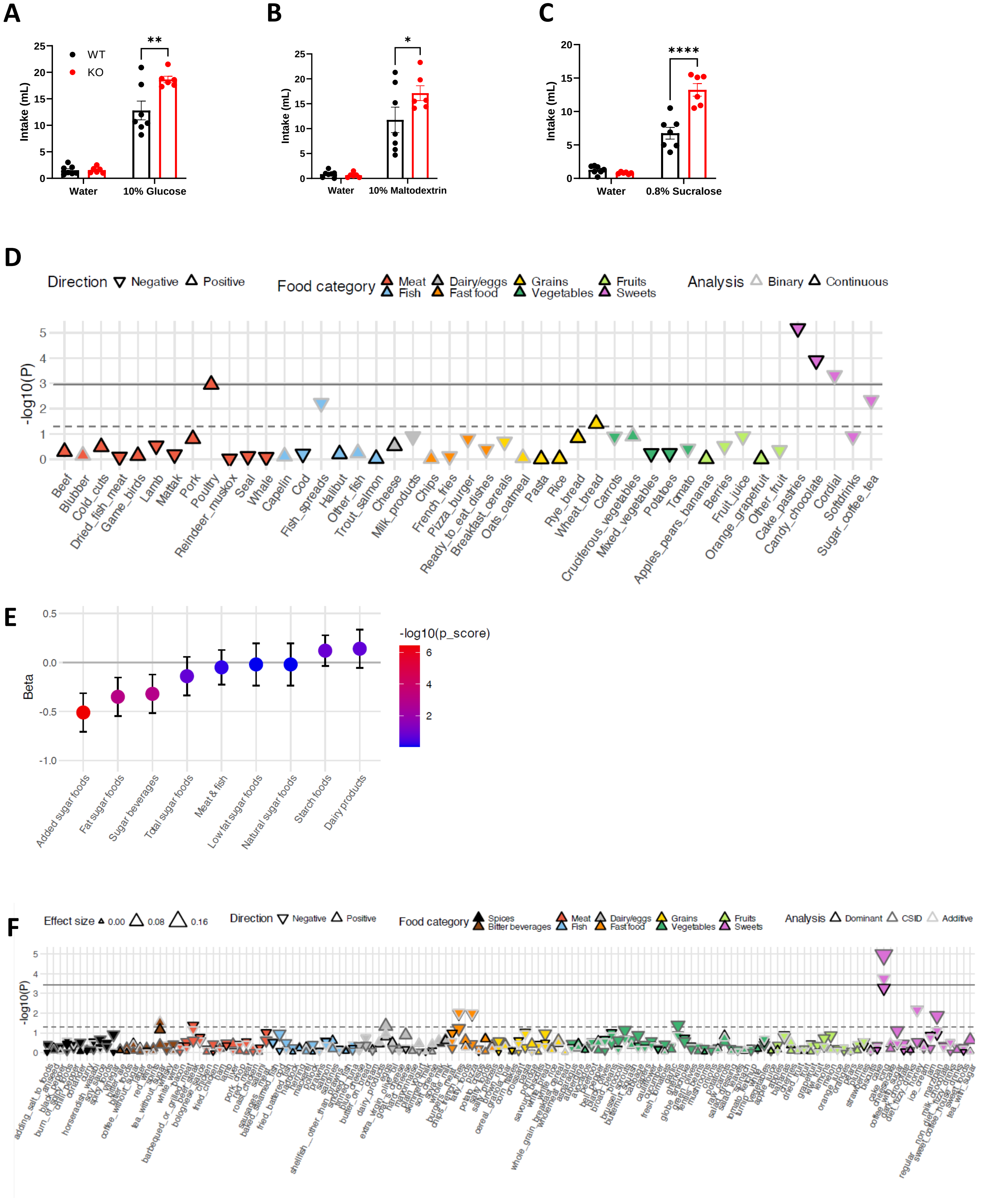
